# Evaluation of computational tools for predicting CRISPR gRNA on-target efficiency in plants

**DOI:** 10.1101/2025.10.15.679654

**Authors:** Zheng Gong, Mengyi Chen, Hui Zhang, Jenny C. Mortimer, José R. Botella

**Author notes:** Co-senior and corresponding authors: Hui Zhang, Jenny Mortimer, José Ramón Botella.

## Abstract

CRISPR technologies has become an integral part of plant biotechnology, synthetic biology and basic plant research, routinely used by researchers for targeted genome modifications. CRISPR guide RNAs (gRNAs) undermines the highly programmable nature of CRISPR, enabling site-specific genome editing. However, different gRNA targets showed highly variable on-target effectiveness and poor gRNA design could amount to wasting valuable scientific resources. There has been broad development of computational and web-based tools for gRNA efficiency predictions but their performances in plant genome editing remains controversial or untested. Hence, in this study, we systematically evaluated over 20 accessible, web-based *in silico* gRNA on-target efficiency prediction tools using an experimental plant genome editing dataset. Excitingly, we identified multiple tools, mostly developed using machine learning, that were highly predictive of gRNA on-target genome editing efficiency *in planta*. The prediction scores assigned to gRNAs in the dataset by these tools were significantly correlated with the frequency of CRISPR-mediated InDels in plants. Furthermore, we evaluated efficiency prediction scores available on popular platforms such as *CRISPOR* and *CRISPR-P* which contain large numbers of non-model plant genomes. Our analysis showed that some prediction scores on *CRISPOR* performed quite well which allows efficient integration of on-target and off-target predictions. Overall, we believe that our study provided insights on improving gRNA design during conventional plant genome editing workflows and should also help unfamiliar researchers interested in CRISPR/SpCas9 genome editing.

## MAIN TEXT

Class II CRISPR systems, such as CRISPR/SpCas9, rely on nucleotide complementarity between the gRNA spacer sequence and genomic DNA sequences to direct target-specific genome editing, enabling unprecedented re-programmability (Jinek et al., 2012, Mao et al., 2019). Consequently, the gRNA also underpins CRISPR on-target efficiency which have shown unpredictable variations for different gRNA targets, particularly in plants (Gong et al., 2025, Slaman et al., 2023, Naim et al., 2020, Moreb and Lynch, 2021). Manual design of effective gRNAs is a challenging endeavour due to the complicated sequence and biochemical factors that have been found to govern gRNA on-target performance (Chen and Wang, 2022, Konstantakos et al., 2022, Moreb and Lynch, 2021). Yet, the functionality of gRNAs is often only assessed in plant regenerants after stable transformation, often requiring laborious tissue culture. Computational tools have been developed to predict gRNA on-target activity. Many of these tools were developed using machine learning (ML) with defined gRNA spacer and structural features. These ML-based efficiency prediction models were trained on experimental datasets comprised of different gRNAs and their quantified genome editing efficiencies as determined in animal cells (Konstantakos et al., 2022, Yan et al., 2018). The selection of feature sets and training experimental datasets have been shown to affect prediction performance and the applicability of ML-based gRNA efficiency prediction tools in plant genome editing has remained controversial (Naim et al., 2020, Slaman et al., 2023). This calls for further investigation into benchmarking and testing these tools to determine the direction of future plant genome editing experiments.

Here, we aimed to systematically evaluate the performance of a comprehensive set of CRISPR/SpCas9 gRNA on-target prediction tools and their relevance in plant applications using a small-scale experimental plant genome editing dataset (Figure 1A). In a past study, we transiently expressed 20 CRISPR/SpCas9 gRNAs in leaves of the dicotyledonous *Nicotiana benthamiana* model plant (Gong et al., 2025). Targeted amplicon sequencing (AmpSeq) was used to quantify genome editing efficiency which was re-analyzed to capture InDel mutations (Figure 1B) (Supplementary Data). Overall, the frequency of CRISPR-mediated InDels observed across the 20 gRNAs varied from around 0% to 30% and was distributed evenly but with some biases, as expected of a small dataset (Figure S1A). This transient expression-based approach was reproducible, with the same gRNA yielding similar results in independent experiments (Figure S1B). Because high-throughput screening is significantly more difficult and less established in plants, we generated a small but relatively robust dataset of gRNAs with *in planta* activity quantified using the accurate and sensitive AmpSeq method. Relatively small gRNA datasets have been used in past studies to evaluate gRNA efficiency prediction scores (Labuhn et al., 2018, Naim et al., 2020, Slamen et al., 2023, Liang et al., 2019, Yan et al., 2018) and to study factors affecting gRNA activity in past studies (Liang et al., 2016).

**Figure 1.**
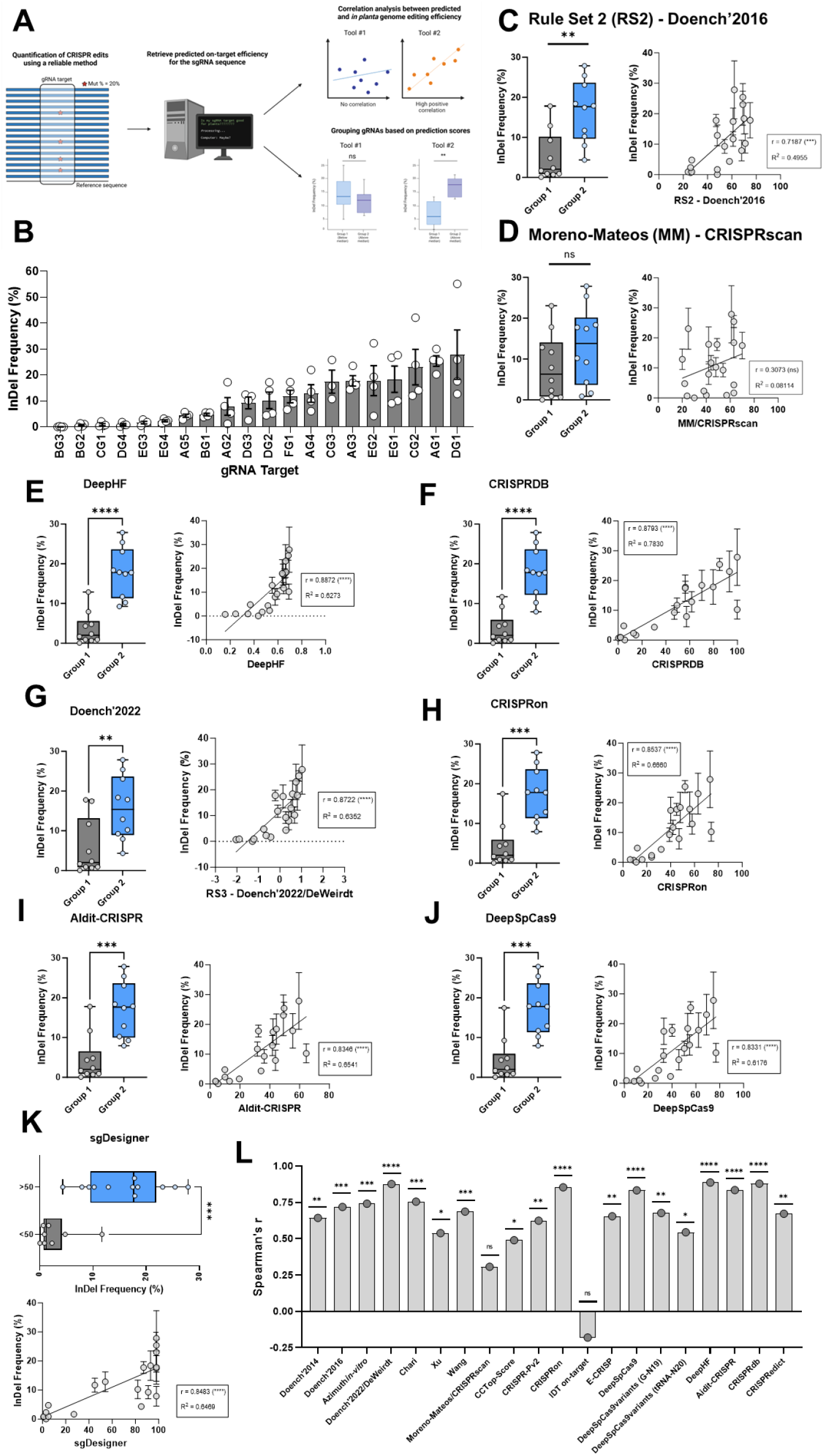
Systematic evaluation of developed gRNA on-target prediction tools. **(A)** Schematic representation of the workflow for this study. An experimental genome editing dataset consists of multiple gRNAs and their on-target genome editing efficiency in plants. The gRNAs were then categorized into groups based on their prediction scores and the overall genome editing efficiencies between the two groups were compared. Linear regression and correlation analyses were also conducted to determine the relationship between gRNA on-target prediction scores and its *in planta* activity. **(B)** Bar graph showing the frequency of reads with CRISPR-mediated InDel mutations across 20 gRNAs as quantified using AmpSeq reported previously and was re-analyzed. Bars represent the mean InDel frequency across biological replicates ± standard error of the mean (SEM). Using this dataset, we evaluated the **(C)** *Doench’2016*, **(D)** *Moreno-Mateos*, **(E)** *DeepHF*, **(F)** *CRISPRDB*, **(G)** *Doench’2022/DeWeirdt*, **(H)** *CRISPRon*, **(I)** *AIdit-CRISPR*, and **(J)** *DeepSpCas9* prediction tools. Box plots (left) representing the CRISPR-mediated InDel frequencies of gRNAs categorized into two groups based on having prediction scores below (group 1) or above (group 2) the median score of the dataset. A parametric unpaired t-test was conducted to determine whether differences in InDel frequencies between the two groups were statistically significant. Linear regression (right) and correlation analyses between the gRNA prediction scores and *in planta* genome editing efficiencies were also conducted and plotted. Spearman’s *r* was calculated as a measure of correlation. **(K)** Evaluating the *sgDesigner* tools for predicting gRNA on-target efficiency in plants. Linear regression and correlation analysis (bottom) of *sgDesigner* score and genome editing efficiency in plants across the 20 gRNAs. Spearman’s *r* was calculated as a measure of correlation. Grouped analysis of *sgDesigner* (top) where gRNAs were categorized into groups with *sgDesigner* scores above or below 50. A parametric unpaired t-test was used to determine whether differences in InDel frequencies between the two groups were statistically significant. **(L)** Summarizing bar graph of Spearman’s correlation of evaluated *in silico* tools with *in planta* genome editing efficiency. The statistical significance of the correlation was marked for each bar. ns, no significance. *, *p*≤0.05. **, *p*≤0.01. ***, *p*≤0.001. ****, *p*≤0.0001.

Next, we queried our set of gRNAs through 21 different *in silico* prediction tools (Table S1) where efficiency scores were retrieved and analyzed. Surprisingly, we found that the popular *Doench’2016* prediction score showed a positive and statistically significant correlation (Spearman’s *r* = 0.7) with genome editing efficiency (Figure 1C), differing from a previous report (Naim et al., 2020). This disagreement may be due to the differences in methods used to quantify genome editing in plants (Figure S3) as further discussed in the Supplementary Notes. We then grouped gRNAs based on their *Doench’2016* scores and found that the group of gRNAs with scores above the median score resulted in significantly higher frequency of InDels compared to the group of gRNAs with lower scores (Figure 1C). By comparison, the *Moreno-Mateos* score could not be used to differentiate groups of gRNAs (Figure 1D). This is consistent with reports in tomato protoplasts using a larger set of gRNAs (Slamen et al., 2023).

Combining correlation and gRNA grouped analysis, we evaluated the remaining 19 prediction tools, many of which were untested in plants. Excitingly, we identified 6 independent tools where the gRNA prediction scores were highly correlated with *in planta* genome editing efficacy, boasting Spearman’s *r* of above 0.8 (Figure 1E – J). These 6 prediction scores also showed different degrees of correlation amongst each other, as expected (Figure S4). Most notably, the prediction score from *CRISPRDB* not only showed a high Spearman’s correlation of nearly 0.88, but the tool was also capable of predicting the range of *in planta* genome editing efficiency across different gRNAs (Figure 1F). This was somewhat observed with *CRISPRon, AIdit-CRISPR* and *DeepSpCas9* tools (Figure 1H – J). Meanwhile, despite showing the highest correlation, a portion of gRNAs were assigned similar *DeepHF* prediction scores and data points clustered together rather than dispersed (Figure 1E). Therefore, we recommend the *CRISPRDB* prediction tool over *DeepHF* as best performing across those evaluated in this study. The majority of top-performing tools were based on deep learning models except *CRISPRDB* and *Doench’2022/DeWeirdt*.

The better predictive performance of *CRISPRDB* may reflect and emphasize the benefit of an ensemble model approach which may be a useful strategy for constructing a tailored gRNA prediction tool by integrating top-performing models in plants (Chen and Wang, 2022). Another interesting tool is *sgDesigner* where its prediction score showed a high correlation (Spearman’s *r* = 0.84) with on-target genome editing efficiency (Figure 1K), but it was not considered top-performing as it assigned either very high or low scores to gRNAs and lacked variance. By splitting gRNAs into groups, we found that the group of gRNAs with *sgDesigner* scores above 50 mediated significantly higher frequency of InDels compared with those below 50 (Figure 1K). All 12 gRNAs with *sgDesigner* scores above 50 mediated detectable levels of InDels. Thus, *sgDesigner* scores could be useful for selecting functional gRNAs but further investigation with a greater number of gRNAs is needed for confirmation. Ultimately, we believe that combining the output and cross-validating predictions across several of these top-performing predictions such as *CRISPRDB, CRISPRon* as well as ones like *sgDesigner* could improve and assist with the design and selection of useful and effective gRNA targets in plant genome editing studies.

*CRISPOR* and *CRISPR-P 2*.*0* are two highly useful gRNA design portals with many non-model plant genomes which are often missing or limited in independent prediction tools. Hence, we evaluated both the *CRISPR-P 2*.*0* on-target and the other 7 prediction scores available on *CRISPOR*, apart from *Doench’2016* and *Moreno-Mateos* (Figure 1L, S5 and S6). The *Chari, Azimuth in-vitro* and *Wang* prediction scores were the most correlative with *in planta* genome editing efficiency (Spearman’s *r* = 0.76, 0.74, and 0.69, respectively) and could serve as useful tools for gRNA design, especially for non-model plants (Figure 1L and S5). The *CRISPR-P v2*.*0* prediction scores were comparably less correlative (Spearman’s *r* = 0.62) and fell short of some other assessed on-target prediction models (Figure 1L and S6).

Overall, we showed that the performance of a broad suite of gRNA on-target prediction tools could be effectively evaluated using an experimental plant genome editing dataset (Figure 1L). As a proof-of-concept, our study successfully found multiple highly accessible prediction tools where their gRNA on-target scores correlated well with *in planta* genome editing efficiency. Effective design of gRNAs using computational tools is crucial to save valuable scientific resources and increase success in plant genome editing. It is notable, however, that our experimental dataset only consisted of 20 gRNAs and these results should also be further validated in other plant species. We call for the support of other researchers with established transient expression-based CRISPR genome editing workflows in specific plant species for simultaneous cross-species validation of our findings or to identify other effective gRNA on-target prediction tools to benefit future genome editing pursuits in plants.

## AUTHOR CONTRIBUTIONS

Z.G., H.Z. J.C.M. and J.R.B. conceptualized and designed the study. Z.G. selected the gRNAs, analyzed and interpreted the data and drafted the manuscript. M.C. contributed to the analysis and interpretation of data. J.C.M., J.R.B. and H.Z. interpreted the data, revised the manuscript and supervised the study. All authors read and approved the final manuscript.

## ACKNOWLEDGMENTS

The authors would like to first acknowledge the UQ Genome Innovation Hub and all its members for helping with the projects, especially Dr. Di Xia and Stacey Anderson. We also acknowledge Yan Zhang, Dr. Karen Massel, and Dr. Peter Crisp for providing valuable advice and suggestions on different aspects of the project. The authors would like to thank all members of the Botella lab at UQ and Zhang lab at SHNU.

## DATA AVAILABILITY STATEMENT

All experimental data were generated and reported in a previous study (Gong et al., 2025). The raw AmpSeq reads are available at Zenodo (DOI: 10.5281/zenodo.15043050) alongside the publication and the analyzed genome editing data are provided in the Supplementary Data. All associated analyses supporting this work have been presented in the Supplementary Data. Any other data or information required to re-analysis will be made available upon request.

