## Supplementary Notes and Information for "Evaluation of computational tools for predicting CRISPR gRNA on-target efficiency in plants"

**AUTHORS & AFFILIATIONS**

Zheng Gong^1, 2, 3^, Mengyi Chen^4^, Hui Zhang^4, *^, Jenny C. Mortimer^1, 2, *^ José R. Botella^3, *^.

^1^School of Agriculture, Food and Wine, Waite Research Institute, The University of Adelaide, Glen Osmond, Australia, 5064.

^2^ARC Centre of Excellence in Plants For Space, The University of Adelaide, Glen Osmond, Australia, 5064.

^3^Plant Genetic Engineering Laboratory, School of Agriculture and Food Sustainability, The University of Queensland, St Lucia, Australia, 4072.

^4^Shanghai Collaborative Innovation Center of Plant Germplasm Resources Development, College of Life Sciences, Shanghai Normal University, Shanghai, China, 200234.

^*^Co-senior and corresponding authors:

Hui Zhang

Jenny Mortimer

José Ramón Botella

This document contains:

Supplementary Notes

Supplementary Methods

Supplementary Figures S1 – S6

Supplementary Table S1

**SUPPLEMENTARY NOTES**

*Analysis of gRNA target sequence for motifs or positional nucleotide preferences*

The 20 gRNA spacers showed widely variable genome editing efficiencies in *N. benthamiana* (Fig 1B), emphasizing the importance of gRNA design and on-target efficiency prediction. Different gRNA sequence motifs and positional nucleotide preferences have been observed in animal systems (Xiang et al., 2021, Xu et al., 2015). Uncovering these features has been crucial for improving gRNA design and could potentially be used to develop on-target efficiency prediction models (Doench et al., 2016, Konstantakos et al., 2022, Wang et al., 2014). We wondered whether efficient gRNAs in our study showed any positional nucleotide preferences. To test this, the 20 gRNAs were split into quartiles based on their *in planta* genome editing efficiency and sequence logos were produced for gRNA spacers in the most and least efficient quartiles (Fig S2). Interestingly, we found slight preferences for thymine nucleotides across the gRNA spacers in the least efficient quartiles. In contrast, gRNA spacers in the most efficient quartiles preferred purines (guanine or adenine) around the middle of the spacer. There is also a frequent occurrence of purines in the nucleotide immediately adjacent to the PAM, known to be critical for gRNA on-target efficiency (Xu et al., 2015, Wang et al., 2014). For inefficient gRNAs, this position has a high prevalence of thymine and, surprisingly, also slightly for guanine. This is generally consistent with preferences identified in larger-scale studies of animal systems (Xiang et al., 2021, Xu et al., 2015, Wang et al., 2014). These findings may provide a starting point for future studies but genome editing datasets from *N. benthamiana* and other plant species with more gRNAs should be analyzed to confirm these observations. Nevertheless, manually designing gRNAs and grasping their on-target efficiency based on known motif or positional nucleotide preferences remain difficult and potentially unreliable. Therefore, developing and benchmarking the performance of gRNA prediction models in plants is a crucial area of future research.

*Importance of genome editing quantification in studying gRNA on-target predictions*

Experimental evaluation of prediction models for gRNA on-target activity in plants was mostly studied as complements to other research in past studies. To our knowledge, there has been a lack of extensive investigation into this topic, even though research in this area has been heavily invested in animal systems (Chari et al., 2015, Chari et al., 2017, Chen and Wang, 2022, Concordet and Haeussler, 2018, Corsi et al., 2023, DeWeirdt et al., 2022, Doench et al., 2016, Doench et al., 2014, Haeussler et al., 2016). Yet, designing suitable, functional and optimal gRNA is absolutely crucial to efficient CRISPR genome editing in plants.

Several studies preceding this work have attempted to evaluate the applicability of gRNA on-target prediction tools for plant genome editing but results have been inconsistent. For example, Naim et al., (2020) transiently expressed 10 gRNAs in *N. benthamiana* to evaluate 8 gRNA efficiency prediction software but found that none were correlative, including the *Doench’2016* prediction score. However, our current work and that reported previously in tomato protoplasts have both found that the *Doench’2016* score may be a predictor of gRNA on-target efficiency in plants. Notably, both latter studies have used the NGS-based AmpSeq to quantify targeted genome editing whereas past studies have relied on reporter transgene knockout assays (Liang et al., 2019, Naim et al., 2020). Corsi et al., (2023) have argued that reporter gene knockout assays are less quantitative and are biased towards specific mutation types. Phenotypic screens to assess gRNA on-target efficiency that rely on the knockout of a functional gene are also dependent on the location in the gene structure that the gRNA targets (Wang et al., 2014). Further emphasis was placed on the use of accurate techniques for quantifying genome edits, such as AmpSeq, when generating experimental datasets for evaluating and training gRNA on-target prediction algorithms (Corsi et al., 2023, Xiang et al., 2021).

The impact of different quantification methods was considered by re-analyzing datasets generated previously (Gong et al., 2025) where we quantified genome edits at these same 20 gRNA targets using 6 other methods: T7E1 analyzed with agarose gel electrophoresis (A), T7E1 analyzed with TapeStation (T), PCR-Capillary Electrophoresis (CE)/Indel Detection by Amplicon Analysis (IDAA), Inference of CRISPR Edits (ICE) analysis of Sanger traces with and without PeakTrace (PT+ or PT-, respectively), and digital droplet PCR (ddPCR). We performed linear regression analyses between the quantified genome editing frequencies and the gRNA’s *Doench’2016* prediction scores (Fig S3). We plotted each technique with the AmpSeq benchmark on the same XY axis. The ddPCR dataset generated a linear regression line that is comparable to AmpSeq with a similar slope and R^2^ (Fig S3F). In contrast, both ICE (PT+) and ICE (PT-) had very different R^2^ and slope in comparison to AmpSeq (Fig S3D, E). To our surprise, T7E1 (A) and T7E1 (T) had closer R^2^ values despite their much lower slope (Fig S3A, B). Overall, we showed that using different techniques to quantify CRISPR genome editing may lead to variable results when evaluating the same gRNA on-target prediction tool. This may explain differences in the outcome of studies and reinstate the importance of selecting a reliable method to assess genome editing efficiency in future work.

**SUPPLEMENTARY METHODS**

*Quantification of genome editing efficiency using targeted amplicon sequencing*

In our previous study (Gong et al., 2025), we transiently expressed 20 individual gRNAs targeting 6 different genes in *Nicotiana benthamiana* and quantified genome editing efficiency using targeted amplicon sequencing (AmpSeq) as described. The raw reads from Illumina MiSeq 150 bp pair end sequencing were re-analyzed using CRISPResso2, accounting strictly for InDels and substitutions were ignored to obtain the InDel frequencies for each sample. This was important because we noted the occurrence of minimal, but detectable levels (<1%) of substitution mutations in some negative control samples. An experimental plant genome editing dataset was produced with 20 gRNAs and AmpSeq-quantified InDel frequencies in 3 to 4 biological replicates each.

*Construction of sequence logo to identify potential positional nucleotide preferences in efficient gRNAs*

The 20 gRNAs were categorized into quartiles based on their on-target genome editing efficiency as measured using AmpSeq. Sequence logos were constructed using WebLogo portal (<https://weblogo.berkeley.edu/logo.cgi>). The spacer sequence and PAM (5’-NGGN-3’) of gRNAs in the most and least efficient quartile were used as input to construct sequence logos.

*Retrieving gRNA on-target efficiency scores from* in silico *prediction tools*

The access portal/platform and website URL for all gRNA prediction tools that were evaluated in this study were provided in Table S1. For the *Doench’2014*, *Doench’2016*, *Chari*, *Xu*, *Wang*, *Moreno-Mateos*, *CCTop*, *Azimuth* *in-vitro* and *WU* scores, we queried the target sequence into the *CRISPOR* web server tool. We selected the SolGenomics *N. benthamiana* genome and the 20 bp, 5’-NGG-3’ PAM sequence for SpCas9. The gRNA sequence and its respective scores were identified and recorded. For the *Doench’2022/DeWeirdt* score, we queried the target sequence on *CRISPick*. No plant genomes were available on *CRISPick*, so we selected the Human GRCh38 as our reference genome for “CRISPRko” analysis using the SpyoCas9 5’-NGG-3’ PAM sequence. Importantly, we used the “Hsu (2013)” tracrRNA for gRNA on-target prediction. The CRISPR-P v2.0 prediction scores were retrieved from the *CRISPR-P v2.0* web server using the *N. benthamiana* genome. The *CRISPRon* prediction scores of gRNAs were retrieved using the zebrafish genome. The *IDT* on-target scores were retrieved from Integrated DNA Technologies’ Custom Alt-R CRISPR-Cas9 guide RNA design tool using the *Homo sapiens* genome. The *E-CRISP* prediction scores were retrieved from the *E-CRISP* web server using the *Arabidopsis thaliana* genome and the “medium” application with “exclude targets with poly T motif” and “exclude targets with poly A motif” turned off as well as the “5’ Preceding Base requirement” changed to “any”. The *DeepSpCas9* and *DeepSpCas9variants* were both retrieved from the *DeepCRISPR* web portal by directly entering the target gene sequence. For *DeepSpCas9variants*, prediction scores for both the SpCas9 “(G/g)N19” and “tRNA-N20” gRNAs were recorded. Notably, our study added a G nucleotide preceding the 20 bp gRNA spacer rather than shortening the sequence to 19 bp as the DeepSpCas9variants “(G/g)N19” setting specifies. The *DeepHF* prediction scores were retrieved from the *DeepHF* web server for SpCas9 gRNAs and with “SpCas9_U6” option selected. The *sgDesigner* and *CRISPRDB* prediction scores were retrieved from the *CRISPRDB* web server. Custom prediction with the “U6 promoter” settings were used. For *CRISPRedict* prediction scores, we used the “Interpret” mode and entered a 30 bp sequence around and including the gRNA target according to the manual. The “U6 promoter” and the “Regression” model were used. Lastly, for the *AIdit-CRISPR* prediction scores, the *Homo sapiens* GRCh38 genome was used with the “*AIdit_ON*” model and SpCas9 was selected as the enzyme. For all these gRNA efficiency predictions, the target sequences were manually entered, and prediction scores were retrieved and recorded.

*Linear regression and correlation analysis between gRNA on-target prediction scores and* in planta *genome editing efficiency*

All linear regression and correlation analyses were conducted using GraphPad Prism 9. For each prediction tool, we performed a correlation analysis to compute Spearman correlation coefficients between the predicted gRNA efficiency score and *in planta* genome editing efficiency, measured as the frequency of InDels using AmpSeq. The correlation is considered statistically significant when the *p-value* < 0.05. The simple linear regression between gRNA on-target prediction scores and its corresponding genome editing efficiencies in plants were computed and plotted using GraphPad Prism 9.

*Grouped analyses by gRNA prediction scores*

Grouped analysis was used to evaluate whether prediction scores can be used to categorize more and less efficient gRNA targets. The 20 gRNAs from the *N. benthamiana* dataset were categorized into one group with higher or one with lower prediction scores relative to the median value. A parametric unpaired *t*-test was used to determine if differences in genome editing efficiencies between the two groups were of statistical significance using GraphPad Prism 9. The same analyses were conducted for all assessed prediction tools unless otherwise stated. A *p*-value < 0.05 was considered statistically significant.

For the *WU* prediction algorithm, we categorized gRNAs with scores of 0 into one group and >0 into another group. A parametric unpaired *t*-test was used to determine if the differences were statistically significant. A *p*-value < 0.05 was considered statistically significant. For *sgDesigner*, we categorized gRNAs with scores <50 into one group and >50 into another group. Similarly, a parametric unpaired *t*-test was used to determine if the differences were statistically significant. A *p*-value < 0.05 was considered statistically significant.

*Comparing CRISPR-mediated mutation quantification methods for producing experimental datasets used to evaluate gRNA on-target efficiency prediction tools*

To investigate the effect of using different genome editing quantification methods to evaluate the performance of prediction models, we retrieved the mutation frequencies for the same sample and gRNA that was quantified using 6 different methods in our previous work (Gong et al., 2025). The CRISPR-mediated mutations in the 20 gRNAs and its samples were quantified using T7E1 separated using agarose gel electrophoresis (A), T7E1 separated using TapeStation (T), PCR-Capillary Electrophoresis (CE) or known as Indel Detection by Amplicon Analysis (IDAA), Sanger sequencing analyzed with Inference of CRISPR Edits (ICE) with and without base calling using Peaktrace (PT+ or PT-, respectively), and lastly with digital droplet PCR (ddPCR). We performed linear regression analysis between the quantified mutation frequency for each method across all gRNAs and its *Doench’2016* prediction score. Linear regression analysis for each technique was performed using GraphPad Prism 9 and was plotted on the same XY axis with that of AmpSeq for comparison. The *R^2^* and linear relationship were also computed.

**SUPPLEMENTARY FIGURES**

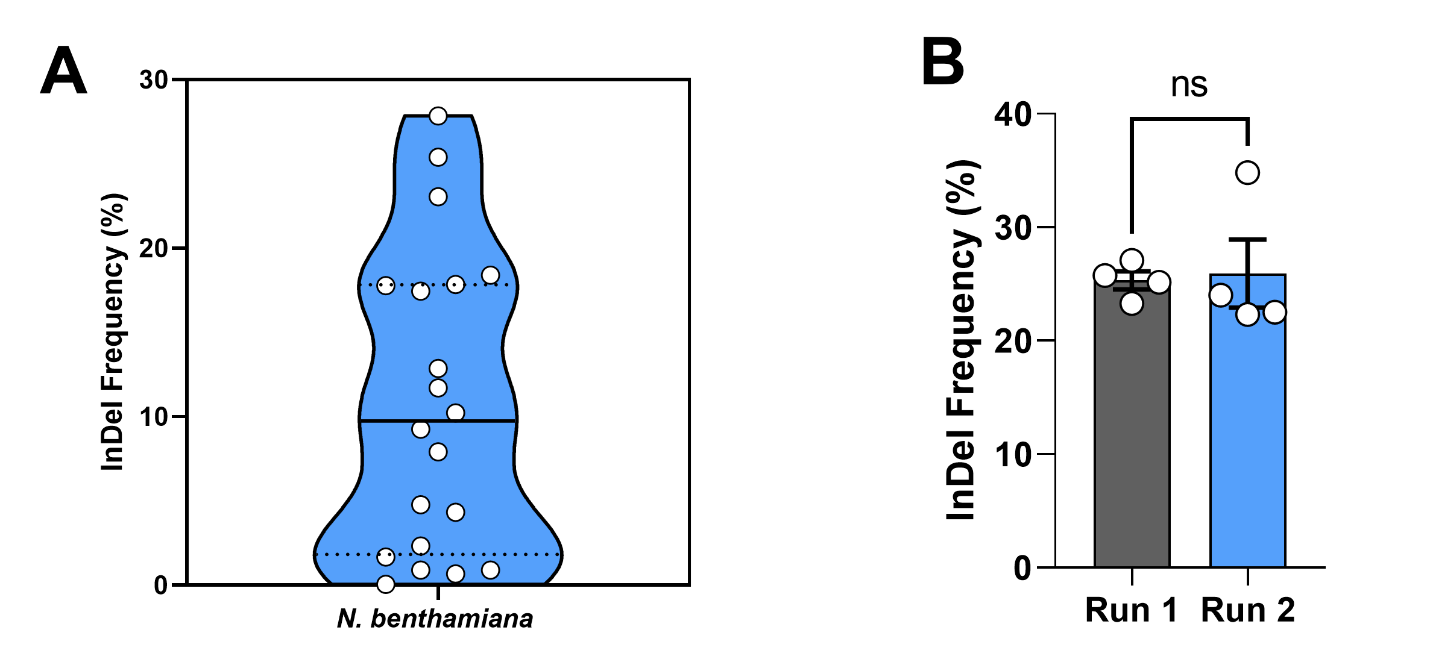

**Figure S1.** Experimental genome editing dataset in *Nicotiana benthamiana*. **(A)** Violin plot showing the distribution of mean InDel frequencies across the 20 gRNAs. The solid line represents the median and the dotted lines represent the two quartiles. **(B)** Genome editing through transient expression of the same gRNA (AG1) across two independent experimental runs. Bars represent the mean InDel frequency ± SEM. The data for Run 1 was displayed in panel A and Run 2 was a second experiment conducted independently. A parametric unpaired *t*-test was conducted to determine if the differences were statistically significant. ns, not significant.

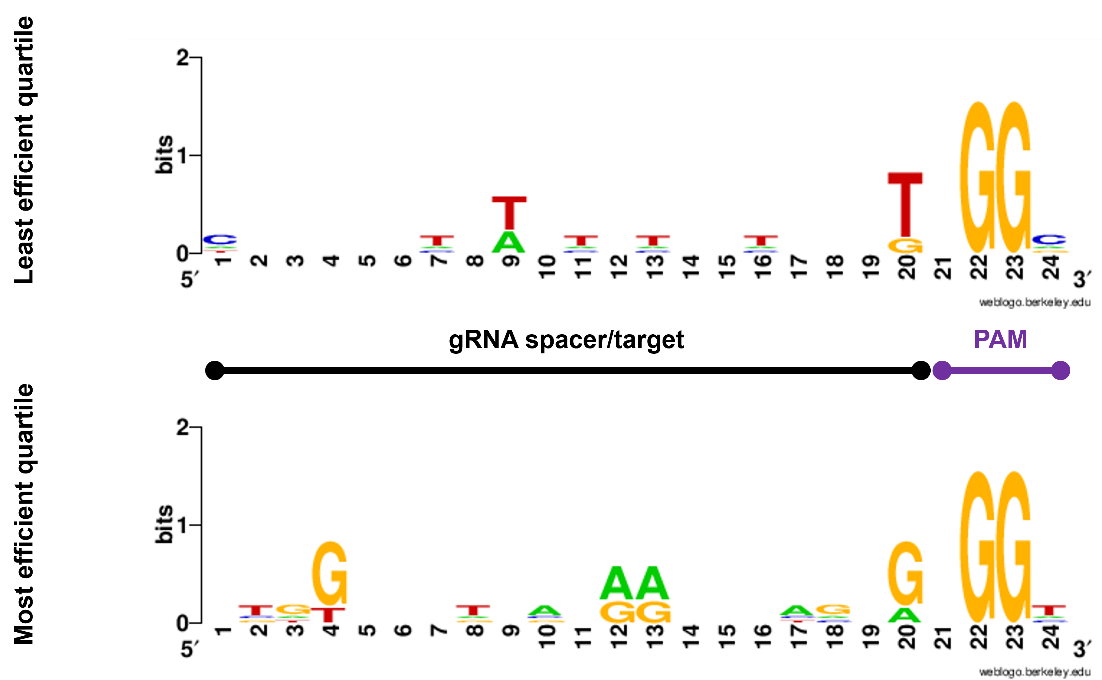

**Figure S2.** Analysis of gRNA sequence features to identify potential positional nucleotide preferences. The genome editing dataset in *N. benthamiana*, consisting of 20 gRNAs was split into quartiles based on their AmpSeq-quantified genome editing efficiency. The gRNA spacer and the PAM sequence of gRNAs in the most and least efficient quartile were inputted into WebLogo to produce sequence logos (Crooks et al. 2004). Bits represent the frequency of nucleotides at each position.

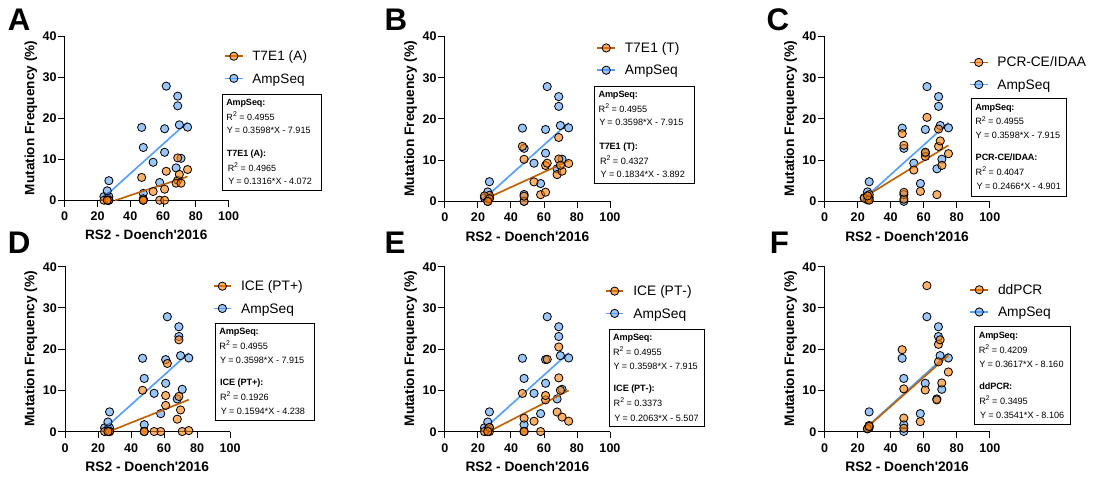

**Figure S3.** The use of different CRISPR genome editing quantification methods to generate the experimental dataset and its effect when evaluating gRNA on-target prediction scores as compared to AmpSeq. Linear regression analysis between genome editing efficiencies of gRNAs quantified using (A) T7E1 (A), (B) T7E1 (T), (C) PCR-CE/IDAA, (D) Sanger sequencing-ICE (PT+), (E) Sanger sequencing-ICE (PT-), (F) ddPCR and the gRNA’s corresponding *Doench’2016* on-target prediction scores were plotted on the same axis as when genome editing efficiencies were quantified using AmpSeq. The R2 and formula of the regression line are shown in the boxes.

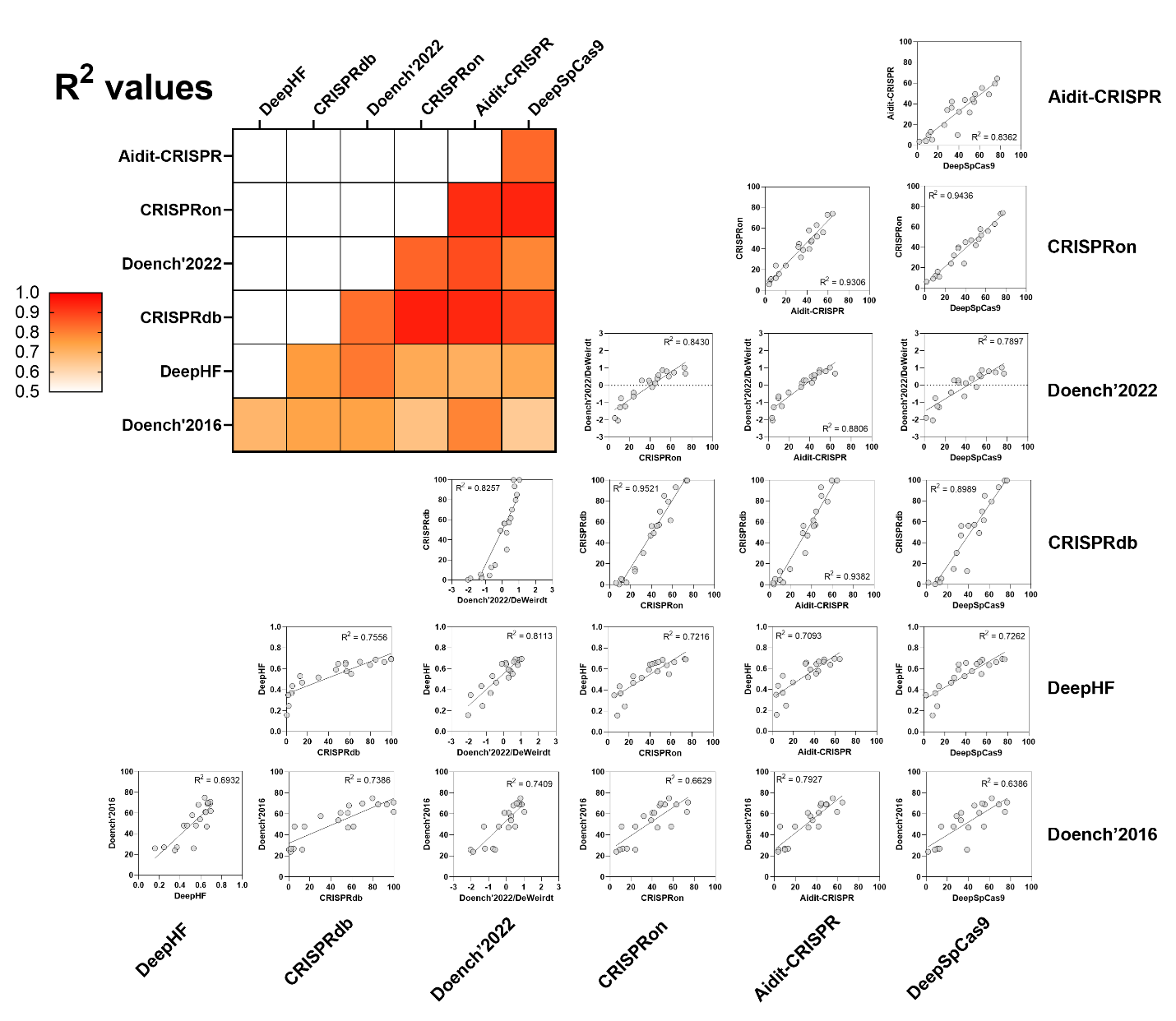

**Figure S4.** Linear regression analysis between the prediction scores of the 6 top-performing gRNA prediction scores. The *Doench’2016* prediction score was also included for comparison. The R^2^ value for the linear regression analyses was represented using a heat map (top left)

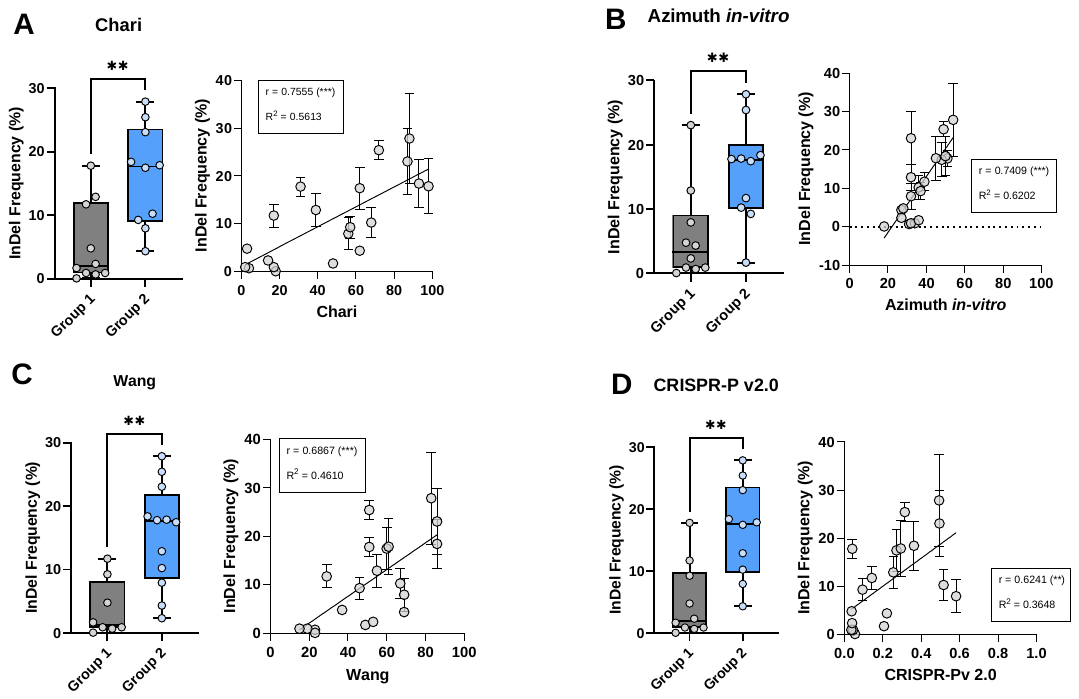

**Figure S5.** Several notable and potentially useful gRNA prediction tools were identified through linear regression and grouped analysis. Analysis of gRNA on-target prediction tools, (A) *Chari*, (B) *Azimuth* *in-vitro*, (C) *Wang* and (D) *CRISPR-P v2.0* are shown. Box and whisker plots for the grouped analysis of each of the prediction scores are shown on the left. The gRNAs were grouped based on their prediction scores. Groups 1 and 2 represents gRNAs with prediction scores lower or higher than the median, respectively. An unpaired, parametric *t*-test was conducted to test for the statistical significance of differences in the mean between the two groups. Linear regression and correlation analysis between *Chari*, *Azimuth* *in-vitro*, *Wang* or *CRISPR-P v2.0* and the gRNAs’ *in planta* genome editing efficiency, as quantified using AmpSeq, was shown on the right of the panels. Each data point represents the mean InDel frequency ± SEM of a gRNA and its corresponding prediction score. The statistical significance of the correlation is shown in brackets. Statistical significance is represented as: ns, no significance. *, *p*≤0.05. **, *p*≤0.01. ***, *p*≤0.001. ****, *p*≤0.0001.

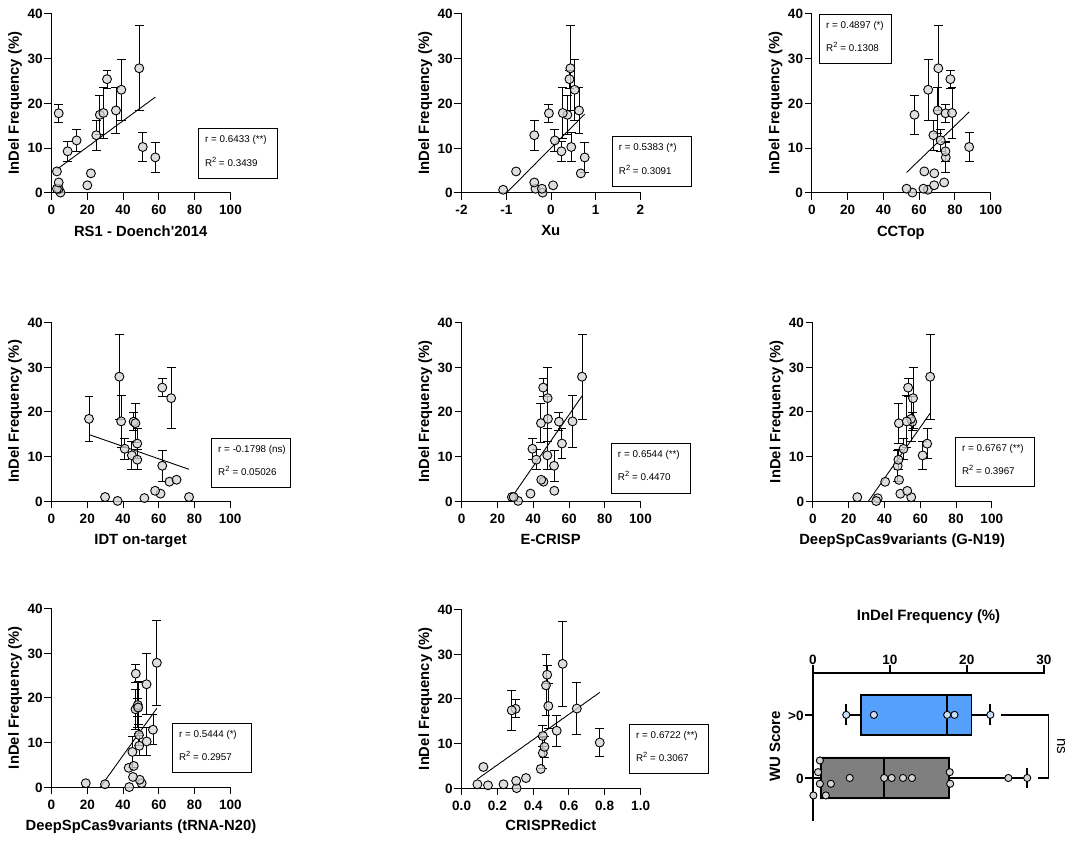

**Figure S6.** Evaluation of remaining gRNA on-target prediction tools. Linear regression and correlation analysis between the gRNA prediction scores for each of the remaining 9 prediction tools and the gRNAs’ *in planta* genome editing efficiency, as quantified using AmpSeq. The statistical significance of the correlation is shown in brackets. For the *WU* prediction tool, gRNAs were grouped based on having scores of 0 (N = 15) or scores greater than 0 (N = 5). An unpaired, parametric *t*-test was conducted to test for the statistical significance of differences in the mean between the two groups. Statistical significance is represented as: ns, no significance. *, *p*≤0.05. **, *p*≤0.01. ***, *p*≤0.001. ****, *p*≤0.0001.

**SUPPLEMENTARY TABLES**

**Table S1.** Table summarizing information for the gRNA prediction tools tested in this study.

| **Prediction tool** | **Access portal** | **Website URL** | **Reference** |
| --- | --- | --- | --- |
| *Doench’2014* | CRISPOR | http://crispor.gi.ucsc.edu/crispor.py | (Doench et al. 2014) |
| *Doench’2016* | CRISPOR | http://crispor.gi.ucsc.edu/crispor.py | (Doench et al. 2016) |
| *Azimuth in-vitro* | CRISPOR | http://crispor.gi.ucsc.edu/crispor.py | CRISPOR |
| *Doench’2022/DeWeirdt* | CRISPick | https://portals.broadinstitute.org/gppx/crispick/public | (DeWeirdt et al. 2022) |
| *Chari* | CRISPOR | http://crispor.gi.ucsc.edu/crispor.py | (Chari et al. 2015) |
| *Xu* | CRISPOR | http://crispor.gi.ucsc.edu/crispor.py | (Xu et al. 2015) |
| *Wang* | CRISPOR | http://crispor.gi.ucsc.edu/crispor.py | (Wang et al. 2014) |
| *Moreno-Mateos* | CRISPOR | http://crispor.gi.ucsc.edu/crispor.py | (Moreno-Mateos et al. 2015) |
| *CCTop* | CRISPOR | http://crispor.gi.ucsc.edu/crispor.py | (Stemmer et al. 2015) |
| *WU* | CRISPOR | http://crispor.gi.ucsc.edu/crispor.py | (Wong et al. 2015) |
| *CRISPR-P v2.0* | CRISPR-P server | http://crispr.hzau.edu.cn/CRISPR2/ | (Lei et al. 2014; Liu et al. 2017) |
| *CRISPRon* | CRISPRon server | https://rth.dk/resources/crispr/crispron/ | (Xiang et al. 2021) |
| *IDT on-target* | IDT server | https://sg.idtdna.com/site/order/designtool/index/CRISPR_CUSTOM | IDT |
| *E-CRISP* | E-CRISP server | http://www.e-crisp.org/E-CRISP/ | (Heigwer et al. 2014) |
| *DeepSpCas9* | DeepCRISPR | https://deepcrispr.info/DeepSpCas9/ | (Kim et al. 2019) |
| *DeepSpCas9variants* | DeepCRISPR | https://deepcrispr.info/DeepSpCas9variants/ | (Kim et al. 2020) |
| *DeepHF* | DeepHF | http://www.deephf.com/#/home | (Wang et al. 2019) |
| *sgDesigner* | CRISPRdb | https://crisprdb.org/wu-crispr-website/ | (Hiranniramol et al. 2020) |
| *CRISPRdb* | CRISPRdb | https://crisprdb.org/custom.html | (Chen and Wang 2022) |
| *CRISPRedict* | CRISPRedict | http://www.crispredict.org/ | (Konstantakos et al. 2022) |
| *AIdit-CRISPR* | AIdit | https://crispr-aidit.com/webServer/gRNA | (Zhang et al. 2023) |
